# Robust estimation of sulcal morphology

**DOI:** 10.1101/452789

**Authors:** Christopher R. Madan

**Affiliations:** School of Psychology, University of Nottingham, Nottingham, United Kingdom

**Keywords:** sulcal width, sulcal depth, age, cortical structure, atrophy, gyrification, cerebral sulci

## Abstract

While it is well established that cortical morphology differs in relation to a variety of inter-individual factors, it is often characterized using estimates of volume, thickness, surface area, or gyrification. Here we developed a computational approach for estimating sulcal width and depth that relies on cortical surface reconstructions output by FreeSurfer. While other approaches for estimating sulcal morphology exist, studies often require the use of multiple brain morphology programs that have been shown to differ in their approaches to localize sulcal landmarks, yielding morphological estimates based on inconsistent boundaries. To demonstrate the approach, sulcal morphology was estimated in three large sample of adults across the lifespan, in relation to aging. A fourth sample is additionally used to estimate test-retest reliability of the approach. This toolbox is now made freely available as supplemental to this paper: https://cmadan.github.io/calcSulc/.

## 1 Introduction

Cortical structure differs between individuals. It is well known that cortical thickness generally decreases with age (Fjell et al., 2009; Hogstrom et al., 2013; Hutton et al., 2009; Lemaitre et al., 2012; Madan & Kensinger, 2016, 2018; Madan, 2018; McKay et al., 2014; Salat et al., 2004; Sowell et al., 2003, 2007); however, a more visually prominent difference is the widening of sulci, sometimes described as “sulcal prominence” (Coffey et al., 1992; Drayer, 1988; Jacoby et al., 1980; Laffey et al., 1984; Tomlinson et al., 1968; Yue et al., 1997). In the literature, this measure has been referred to using a variety of names, including sulcal width, span, dilation, and enlargement, as well as fold opening. With respect to aging and brain morphology, sulcal width has been assessed qualitatively by clinicians as an index of cortical atrophy (Coffey et al., 1992; Drayer, 1988; Laffey et al., 1984; Pasquier et al., 1996; Scheltens et al., 1997; Tomlinson et al., 1968). An illustration of age-related differences in sulcal morphology is shown in Figure 1.

**Figure 1.**
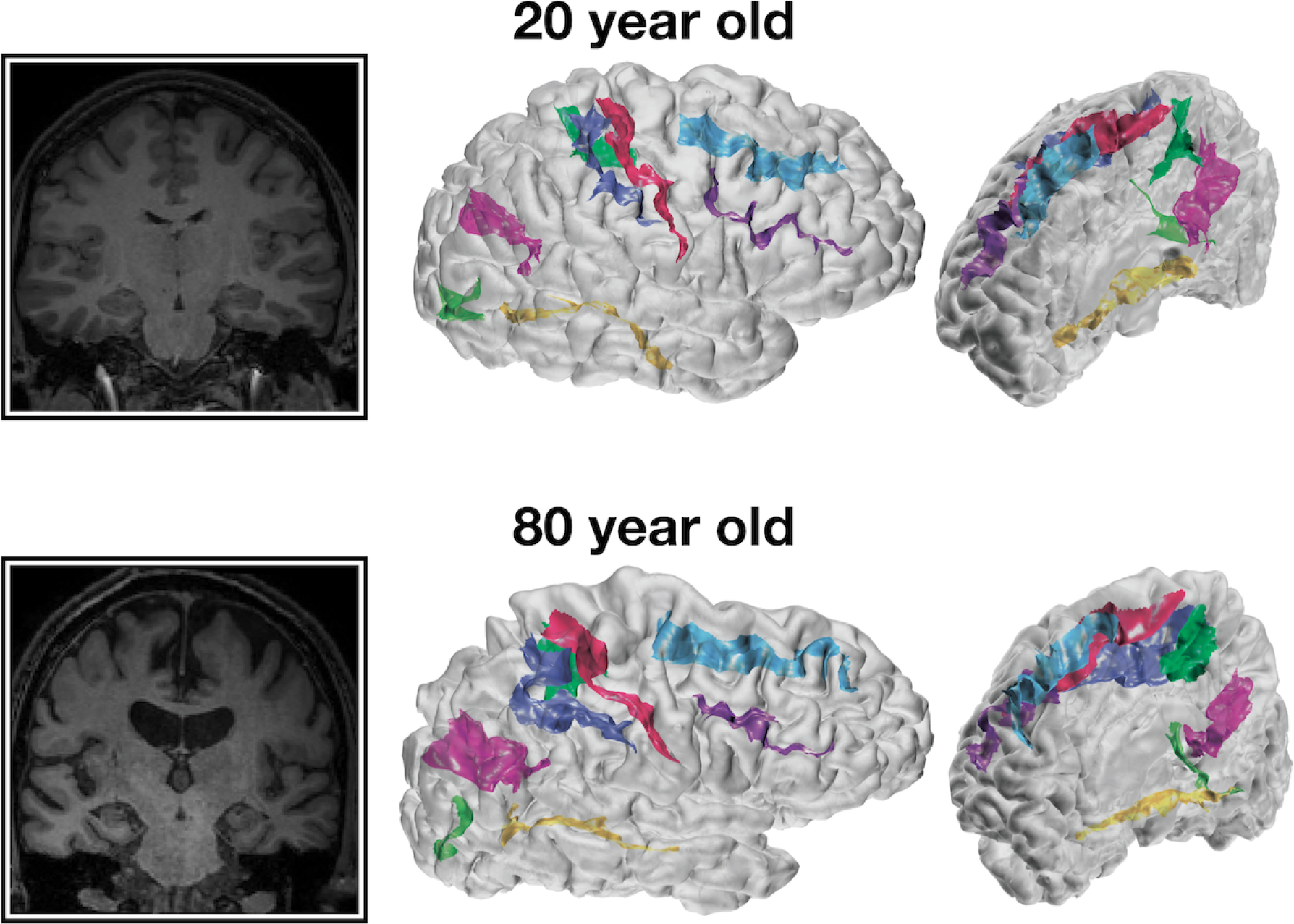
Representative coronal slices and cortical surfaces with sulcal identification for 20- and 80-year-old individuals.

Using quantitative approaches, sulcal width has been shown to increase with age (Kochunov et al., 2005, 2008; Liu et al., 2010, 2013) likely relating to subsequent findings of age-related decreases in cortical gyrification (Cao et al., 2017; Hogstrom et al., 2013; Madan & Kensinger, 2016, 2018; Madan, 2018). Sulcal widening has also been shown to be associated with decreases in cognitive abilities (Liu et al., 2011) and physical activity (Lamont et al., 2014). With respect to clinical conditions, increased sulcal width has been found in dementia patients relative to healthy controls (Andersen et al., 2015; Hamelin et al., 2015; Huckman et al., 1975; Liu et al., 2012; Ming et al., 2015; Plocharski & Østergaard, 2016; Reiner et al., 2012), as well as with schizophrenia patients (Largen et al., 1984; Palaniyappan et al., 2015; Rieder et al., 1979) and mood disorders (Elkis et al., 1995).

One of the most common programs for conducting cortical surface analyses is FreeSurfer (Fischl, 2012). Unfortunately, though FreeSurfer reconstructs cortical surfaces, it does not estimate sulcal width or depth, leading researchers to use FreeSurfer along with another surface analysis program, BrainVISA (Kochunov et al., 2012; Mangin, Rivière, et al., 2004; Mangin, Riviere, et al., 2004; Rivière et al., 2002), to characterize cortical thickness along with sulcal morphology (e.g, Cai et al., 2017; Lamont et al., 2014; Liu et al., 2011, 2013; Pizzagalli et al., 2017). While this combination allows for the estimation of sulcal morphology in addition to standard measures such as cortical thickness, FreeSurfer and BrainVISA rely on different anatomical landmarks (Mikhael et al., 2018) which can yield differences in their resulting cortical surface reconstructions (Lee et al., 2006). Admittedly, determining the boundaries for an individual sulcus and incorporating individual cortical variability is difficult (Campero et al., 2014; John et al., 2006; Mikhael et al., 2018; Ono et al., 1990; Rhoton, 2007; ten Donkelaar et al., 2018; Welker, 1990). While an ennumerate amount of other methods have already been proposed to identify and characterize sulcal morphology (e.g., Andreasen et al., 1994; Auzias et al., 2015; Beeston & Taylor, 2000; Behnke et al., 2003; Eskildsen et al., 2005; Im et al., 2010; Jones et al., 2000; Le Goualher et al., 1996, 1998; Le Troter et al., 2012; Li & Shen, 2011; Li et al., 2010; Lohmann & von Cramon, 2000; Lohmann et al., 2008; Nowinski et al., 1996; Oguz et al., 2008; Perrot et al., 2011; Royackkers et al., 1999; Thompson et al., 1996; Vaillant & Davatzikos, 1997; Yang & Kruggel, 2008; Yun et al., 2013), ultimately these all are again using different landmarks than FreeSurfer uses for cortical parcellations (i.e., volume, thickness, surface area, gyrification). Note that, though FreeSurfer itself does compute sulcal maps, these are computed as normalized depths, not in real-world units (e.g. Kippenhan et al., 2005), furthermore, these are also independent of sulcal width information.

Here we describe a procedure for estimating sulcal morphology and report age-related differences in sulcal width and depth using three large samples of adults across the lifespan: two of these datasets are from Western samples, Dallas Lifespan Brain Study (DLBS) and Open Access Series of Imaging Studies (OASIS), as well as one East Asian sample, Southwest University Adult Lifespan (SALD), as potential differences between populations have been relatively understudied (Leong et al., 2017; Madan, 2017). To further validate the method, test-retest reliability was also assessed using a sample of young adults who were scanned ten times within the span of a month (Chen et al., 2015; Madan & Kensinger, 2017b). All four of these datasets are open-access and have sufficient sample sizes to be suitable for brain morphology research (Madan, 2017). This procedure has been implemented as a MATLAB toolbox, `calcSulc`, that calculates sulcal morphology–both width and depth–using files generated as part of the standard FreeSurfer cortical reconstruction and parcellation pipeline. This toolbox is now made freely available as supplemental to this paper: https://cmadan.github.io/calcSulc/.

## 2 Materials and Methods

### 2.1 Datasets

#### OASIS

This dataset consisted of 314 healthy adults (196 females), aged 18–94 (see Figure 2), from the Open Access Series of Imaging Studies (OASIS) cross-sectional dataset (http://www.oasis-brains.org) (Marcus et al., 2007). Participants were recruited from a database of individuals who had (a) previously participated in MRI studies at Washington University, (b) were part of the Washington University Community, or (c) were from the longitudinal pool of the Washington University Alzheimer Disease Research Center. Participants were screened for neurological and psychiatric issues; the Mini-Mental State Examination (MMSE) and Clinical Dementia Rating (CDR) were administered to participants aged 60 and older. To only include healthy adults, participants with a CDR above zero were excluded; all remaining participants scored 25 or above on the MMSE. Multiple T1 volumes were acquired using a Siemens Vision 1.5 T with a MPRAGE sequence; only the first volume was used here. Scan parameters were: TR=9.7 ms; TE=4.0 ms; flip angle=10°; voxel size=1.25×1×1 mm. Age-related comparisons for volumetric and fractal dimensionality measures from the OASIS dataset were previously reported in Madan and Kensinger (2017a), Madan and Kensinger (2018), and Madan (2019) ^1^.

**Figure 2.**
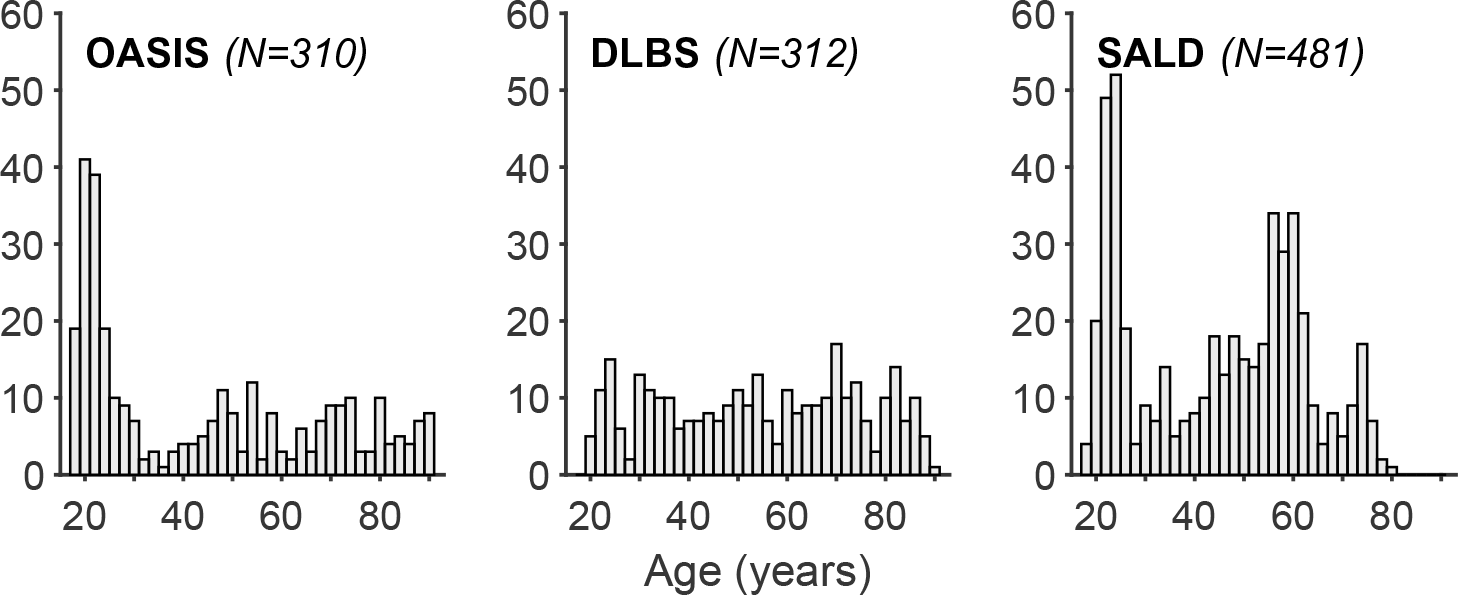
Histogram of age distribution for the three aging datasets: OASIS, DLBS, and SALD, only for participants included in the sulcal morphology analyses. Each bar corresponds to a two-year age-range bin.

#### DLBS

This dataset consisted of 315 healthy adults (198 females), aged 20–89 (see Figure 2), from wave 1 of the Dallas Lifespan Brain Study (DLBS), made available through the International Neuroimaging Data-sharing Initiative (INDI) (Mennes et al., 2013) and hosted on the Neuroimaging Informatics Tools and Resources Clearinghouse (NITRC) (Kennedy et al., 2016) (http://fcon_1000.projects.nitrc.org/indi/retro/dlbs.html). Participants were screened for neurological and psychiatric issues. No participants in this dataset were excluded *a priori*. All participants scored 26 or above on the MMSE. T1 volumes were acquired using a Philips Achieva 3 T with a MPRAGE sequence. Scan parameters were: TR=8.1 ms; TE=3.7 ms; flip angle=12°; voxel size=1×1×1 mm. See Kennedy et al. (2015) and Chan et al. (2014) for further details about the dataset. Age-related comparisons for volumetric and fractal dimensionality measures from the DLBS dataset were previously reported in Madan and Kensinger (2017a), Madan and Kensinger (2018), and Madan (2019) ^1^.

#### SALD

This dataset consisted of 483 healthy adults (303 females), aged 19–80 (see Figure 2), from the Southwest University Adult Lifespan Dataset (SALD) (Wei et al., 2018), also made available through INDI and hosted on NITRC (http://fcon_1000.projects.nitrc.org/indi/retro/sald.html). No participants in this dataset were excluded *a priori*. T1 volumes were acquired using a Siemens Trio 3 T with a MPRAGE sequence. Scan parameters were: TR=1.9 s; TE=2.52 ms; flip angle=9°; voxel size=1×1×1 mm.

#### CCBD

This dataset consisted of 30 healthy adults (15 females), aged 20–30, from the Center for Cognition and Brain Disorders (CCBD) at Hangzhou Normal University (Chen et al., 2015). Each participant was scanned for 10 sessions, occurring 2-3 days apart over a one-month period. No participants in this dataset were excluded *a priori*. T1 volumes were acquired using a SCANNER with a FSPGR sequence. Scan parameters were: TR=8.06 ms; TE=3.1 ms; flip angle=8°; voxel size: 1×1×1 mm. This dataset is included as part of the Consortium for Reliability and Reproducibility (CoRR) (Zuo et al., 2014) as HNU1. Test-retest comparisons for volumetric and fractal dimensionality measures from the CCBD dataset were previously reported in Madan and Kensinger (2017b)^1^.

### 2.2 Procedure

Data were analyzed using FreeSurfer v6.0 (https://surfer.nmr.mgh.harvard.edu) on a machine running Red Hat Enterprise Linux (RHEL) v7.4. FreeSurfer was used to automatically volumetrically segment and parcellate cortical and subcortical structures from the T1-weighted images (Fischl, 2012; Fischl & Dale, 2000) FreeSurfer’s standard pipeline was used (i.e., `recon-all`). No manual edits were made to the surface meshes, but surfaces were visually inspected. Cortical thickness is calculated as the distance between the white matter surface (white-gray interface) and pial surface (gray-CSF interface). Gyrification was also calculated using FreeSurfer, as described in Schaer et al. (2012). Cortical regions were parcellated based on the Destrieux et al. (2010) atlas, also part of the standard FreeSurfer analysis pipeline.

## 3 Calculation

Here we outline a novel, simple yet robust, automated approach for estimating sulcal width and depth, based on intermediate files generated as part of the standard FreeSurfer analysis pipeline. This procedure and functionality has been implemented in an accompanying MATLAB toolbox, `calcSulc`. The toolbox is supplemental material to this paper and is made freely available: https://cmadan.github.io/calcSulc/.

For each individual sulcus (for each hemisphere and participant), the following approach was used to characterize the sulcal morphology. The procedure has been validated and is supported for the following sulci: central, post-central, superior frontal, inferior frontal, parieto-occipital, occipito-temporal, middle occipital and lunate, and marginal part of the cingulate (`S_central`, `S_postcentral`, `S_front_sup`, `S_front_inf`, `S_parieto_occipital`, `S_oc-temp_med&Lingual`, `S_oc_middle&Lunatus`, `S_cingul-Marginalis`). All of the sulci are labeled in Figure 3.

**Figure 3.**
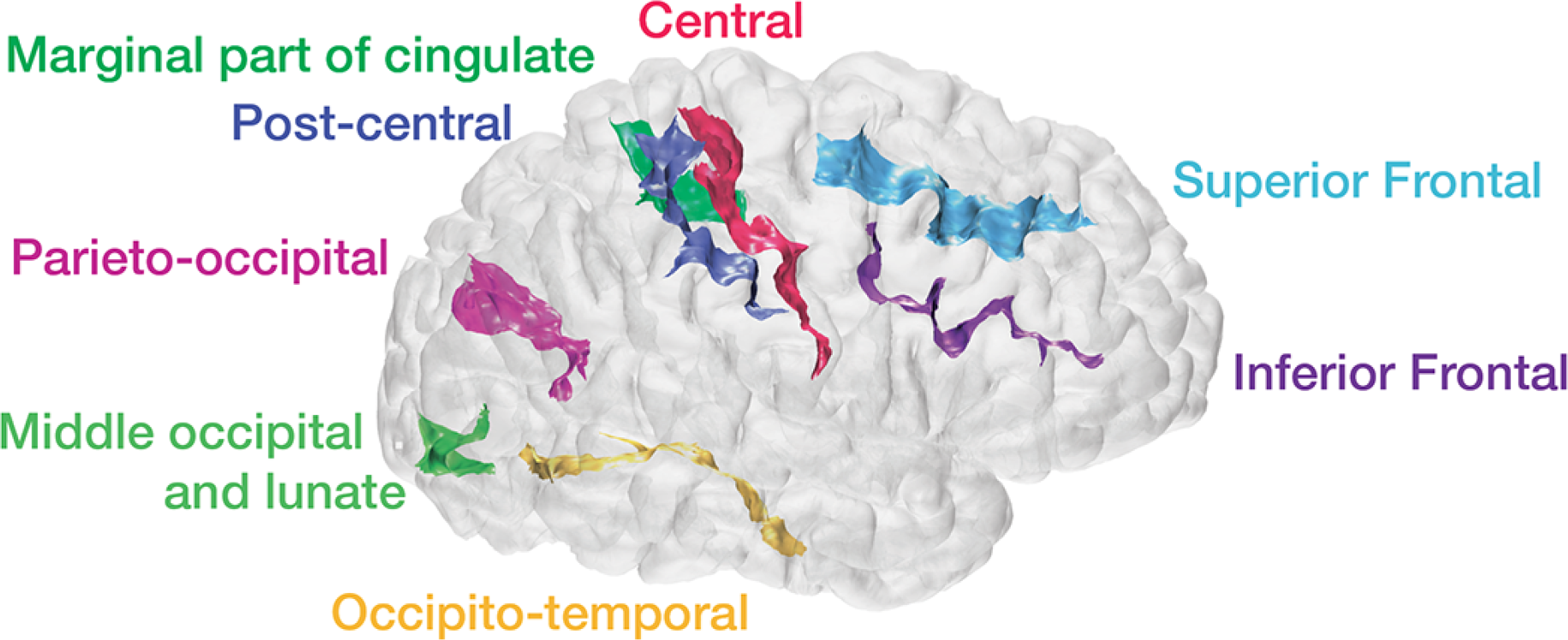
Example cortical surface with estimated sulci identified and labelled.

**Figure 4.**
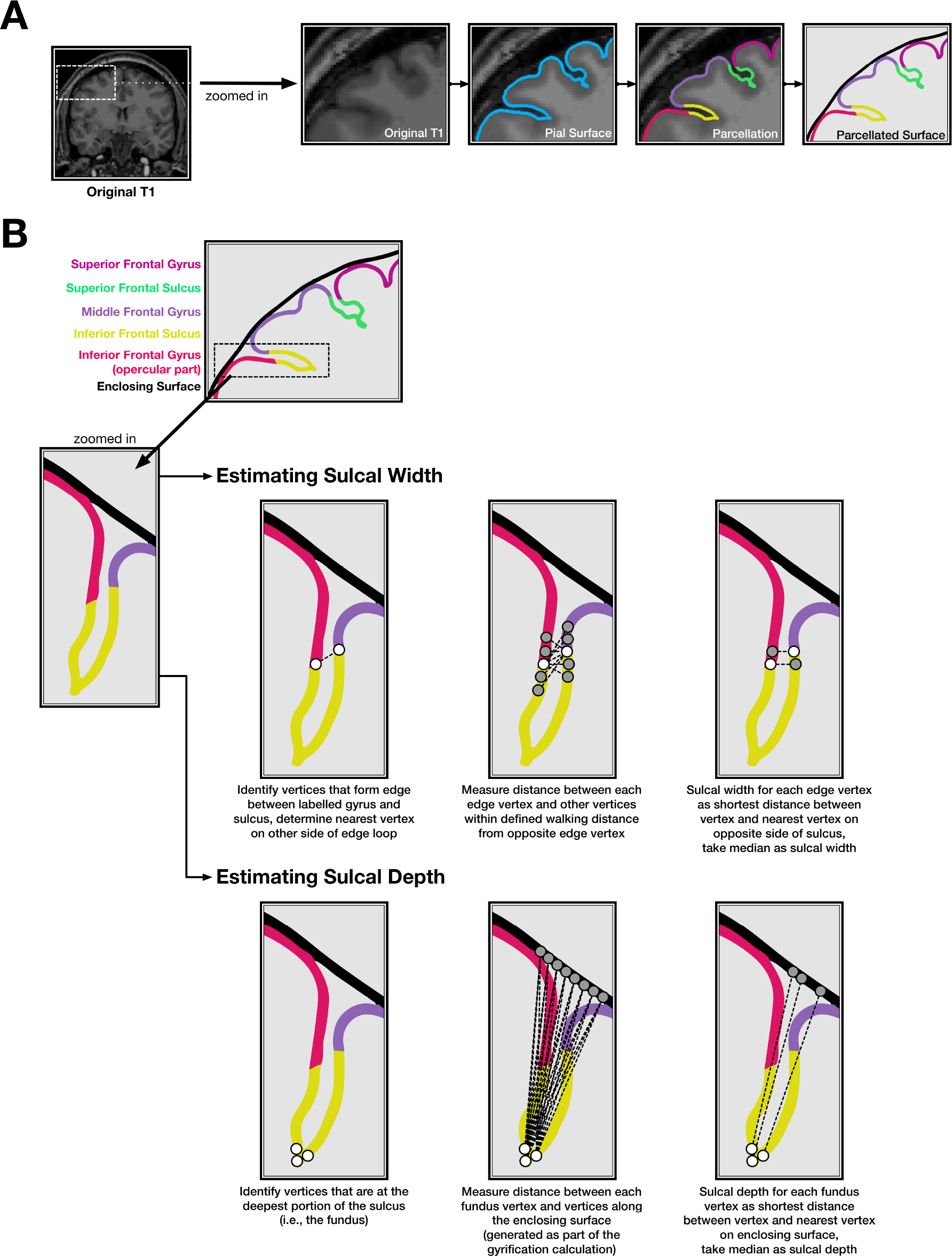
Illustration of the sulcal morphology method. (A) Cortical surface estimation and sulcal identification, as output from FreeSurfer. (B) Sulcal width and depth estimation procedure. Note that the surface mesh and estimation algorithm use many more vertices than shown here.

First the pial surface and Destrieux et al. (2010) parcellation labels were read into MATLAB by using the FreeSurfer-MATLAB toolbox provided alongside FreeSurfer (`calcSulc_load`), this consists of the `?h.pial` (FreeSurfer cortical surface mesh) `?h.aparc.a2009s.annot` (FreeSurfer parcellation annotation) files. Using this, the faces associated with the individual sulcus were isolated as a 3D mesh (`calcSulc_isolate`).

The width of each sulcus (`calcSulc_width`) was calculated by determining which vertices lay on the boundary of the sulcus and the adjacent gyrus. An iterative procedure was then used to determine the ‘chain’ of edges that would form a contiguous edge-loop that encircle the sulcal region (`calcSulc_getEdgeLoop`). This provided an exhaustive list of all vertices that were mid-way between the peak of the respective adjacent gyri and depth of the sulcus itself. For each vertex in this edge-loop, the nearest point in 3D space that was *not* neighbouring in the loop was determined, with the goal of finding the nearest vertex in the edge that was on the opposite side of the sulcus–i.e., a line between these two vertices would ‘bridge’ across the sulcus. Since these nearest vertices in the edge loop are not necessarily the nearest vertex along the opposite sulcus wall, an exhaustive search (walk) was performed, moving up to a 4 edges from the initially determined nearest vertex (configurable as `options.setWidthWalk`). The sulcal width was then taken as the median of these distances that bridged across the sulcus.

The depth of each sulcus (`calcSulc_depth`) additionally used FreeSurfer’s sulcal maps (`?h.sulc`) to determine the relative inflections in the surface mesh, which would be in alignment with the gyral crown. The deepest points of the sulcus, i.e., the sulcal fundus, were taken as the 100 vertices within the sulcus with the lowest values in the sulcal map. For these 100 vertices, the shortest (i.e., Euclidean) distance to the smoothed enclosing surface was calculated (generated by FreeSurfer’s built-in gyrification analysis [`?h.pial-outer-smoothed`], Schaer et al., 2012), and the median of these was then taken as the sulcal depth. While the use of a Euclidean distance here underestimates the true sulcal depth, it is nonetheless robust (as demonstrated in the present work) and does not markedly differ from other algorithmic approaches for estimating sulcal depth for much of the cortex (see Yun et al., 2013, for a comparison).

Sulcal morphology, width and depth, was estimated for eight major sulci in each hemisphere: central, post-central, superior frontal, inferior frontal, parieto-occipital, occipito-temporal, middle occipital and lunate, and marginal part of the cingulate (`S_central`, `S_postcentral`, `S_front_sup`, `S_front_inf`, `S_parieto_occipital`, `S_oc-temp_med&Lingual`, `S_oc_middle&Lunatus`, `S_cingul-Marginalis`). Preliminary analyses additionally included superior and inferior temporal sulci and intraparietal sulcus but these were removed from further analysis when the sulci width estimation was found to fail to determine a closed boundary edge-loop at an unacceptable rate (*>* 10%) for at least one hemisphere. This edge boundary determination failed when parcellated regions were labeled by FreeSurfer to comprise at least two discontinuous regions, such that they could not be identified using a single edge loop. Nonetheless, sulcal measures failed to be estimated for some participants, resulting in final samples of 310 adults from the OASIS dataset, 312 adults from the DLBS dataset, 481 adults from the SALD dataset, and 30 adults from the CCBD dataset (see Figure 2).

### 3.1 Test-retest reliability

Test-retest reliability was assessed as intraclass correlation coefficient (*ICC*), which can be used to quantify the relationship between multiple measurements (Asendorpf & Wallbott, 1979; Bartko, 1966; Chen et al., 2018; Hallgren, 2012; Koo & Li, 2016; Madan & Kensinger, 2017b; Rajaratnam, 1960; Shrout & Fleiss, 1979). McGraw and Wong (1996) provide a comprehensive review of the various *ICC* formulas and their applicability to different research questions. *ICC* was calculated as the one-way random effects model for the consistency of single measurements, i.e., *ICC*(1, 1). As a general guideline, *ICC* values between .75 and 1.00 are considered ‘excellent,’ .60–.74 is ‘good,’ .40–.59 is ‘fair,’ and below .40 is ‘poor’ (Cicchetti, 1994).

## 4 Results & Discussion

### 4.1 Age-related differences in sulcal morphology

Scatter plots showing the relationships between each individual sulcal width and depth and age, for the OASIS dataset, are shown in Figure 5; the corresponding correlations for all datasets are shown in Tables 1 and 2. The width and depth of the central and post-central sulci appear to be particularly correlated with age, with wider and shallower sulci in older adults. Age-related differences in sulcal width and depth and generally present in other sulci as well, but are generally weaker.

**Figure 5.**
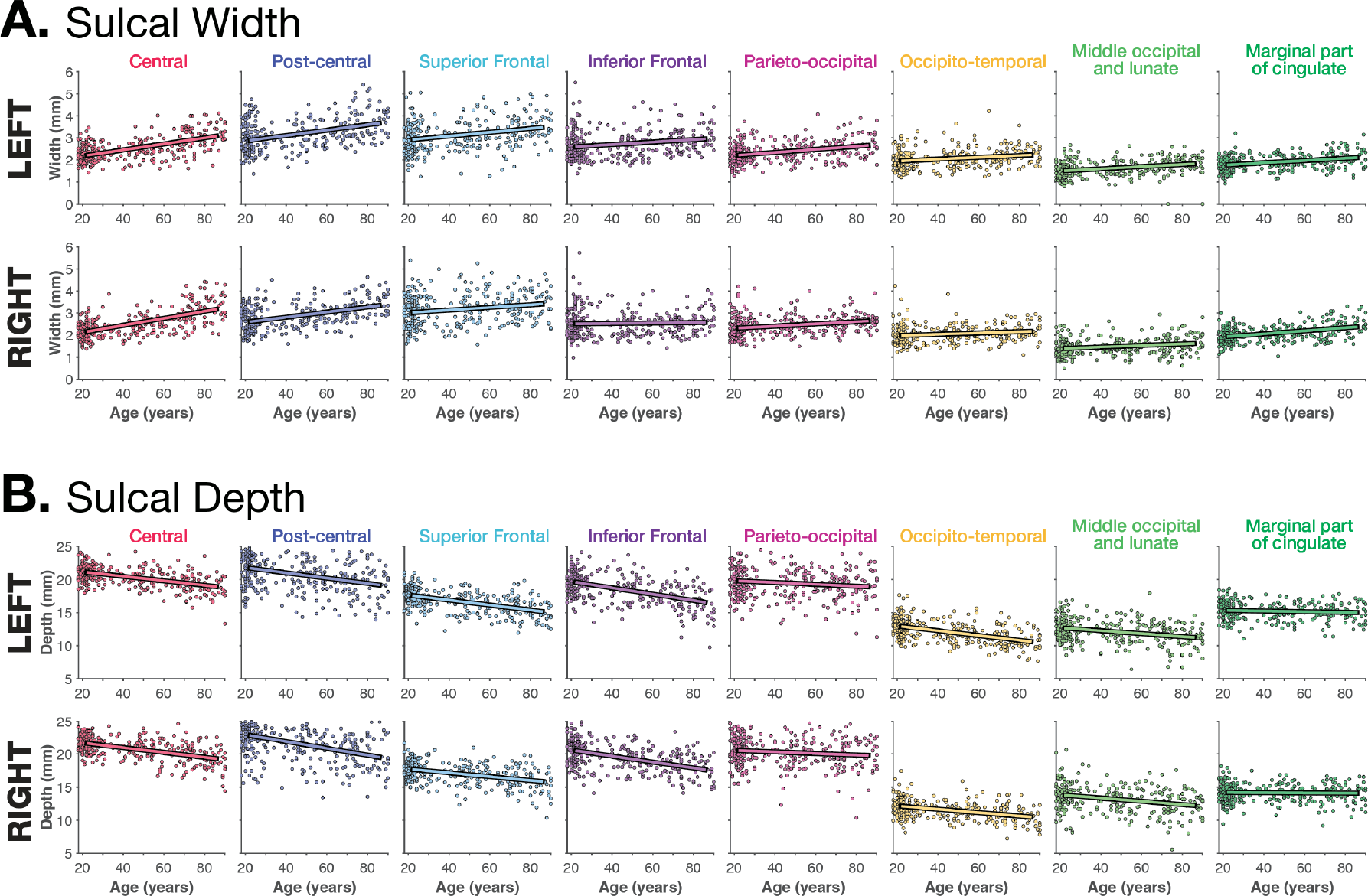
Relationship between (A) sulcal depth and (B) width for each of the sulci examined, based on the OASIS dataset.

**Table 1.** Correlations between sulcal width and age for each sulci and hemisphere, for each of the three lifespan datasets examined. Test-retest reliability, ICC(1, 1), is also included from the CCBD dataset. ^†^FreeSurfer labels in version 6.0; labels are named slightly different in version 5.3. ICC values between .75 and 1.00 are considered ‘excellent,’ .60–.74 is ‘good,’ .40–.59 is ‘fair,’ and below .40 is ‘poor’ (Cicchetti, 1994).

| Sulci Name | FreeSurfer Label† | Hemi. | OASIS | DLBS | SALD | CCBD |  |
| --- | --- | --- | --- | --- | --- | --- | --- |
|  |  |  | *r(Age)* | *r(Age)* | *r(Age)* | *ICC(1,1)* | 95% CI of ICC |
| Central | S_central | L | .586 | .486 | .322 | .858 | [ 0.785, 0.918] |
|  |  | R | .632 | .523 | .294 | .842 | [ 0.764, 0.908] |
| Post-central | S_postcentral | L | .413 | .391 | .198 | .764 | [ 0.660, 0.858] |
|  |  | R | .460 | .436 | .213 | .864 | [ 0.794, 0.922] |
| Superior Frontal | S_front_sup | L | .281 | .421 | .055 | .797 | [ 0.703, 0.880] |
|  |  | R | .205 | .291 | .035 | .843 | [ 0.764, 0.909] |
| Inferior Frontal | S_front_inf | L | .217 | .323 | -.037 | .775 | [ 0.675, 0.865] |
|  |  | R | .043 | .222 | -.036 | .831 | [ 0.748, 0.901] |
| Parieto-occipital | S_parieto_occipital | L | .348 | .279 | .145 | .616 | [ 0.486, 0.753] |
|  |  | R | .257 | .357 | .213 | .682 | [ 0.561, 0.802] |
| Occipito-temporal | S_oc-temp_med&Lingual | L | .227 | .270 | -.055 | .660 | [ 0.535, 0.786] |
|  |  | R | .168 | .189 | .017 | .692 | [ 0.572, 0.808] |
| Middle occipital and lunate | S_oc_middle&Lunatus | L | .306 | .271 | .145 | .605 | [ 0.474, 0.744] |
|  |  | R | .212 | .177 | .023 | .625 | [ 0.496, 0.760] |
| Marginal part of cingulate | S_cingul-Marginalis | L | .340 | .275 | .075 | .783 | [ 0.685, 0.871] |
|  |  | R | .430 | .382 | .161 | .757 | [ 0.651, 0.853] |
| Mean |  |  | .636 | .592 | .227 | .907 | [ 0.856, 0.947] |

**Table 2.**
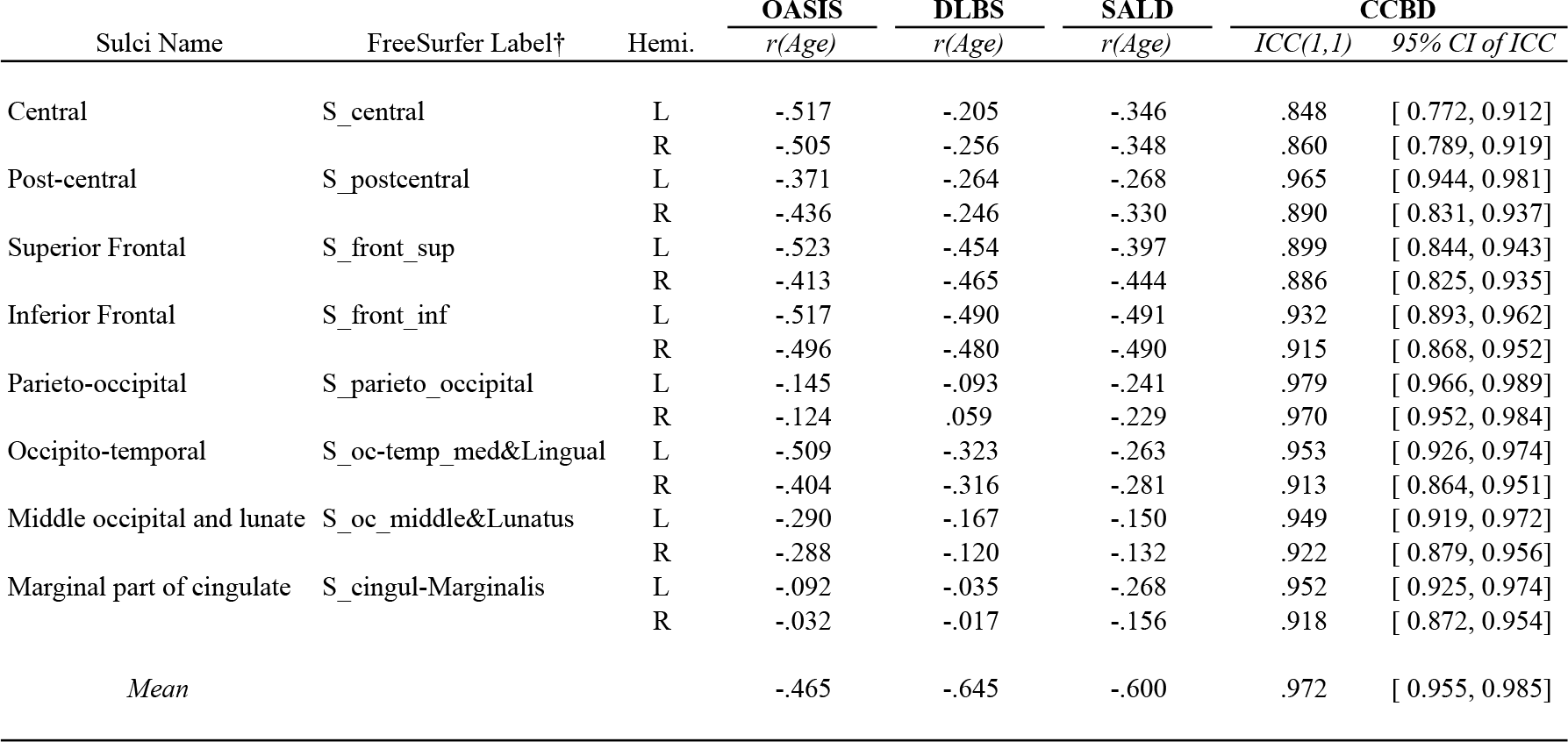
Correlations between sulcal depth and age for each sulci and hemisphere, for each of the three lifespan datasets examined. Test-retest reliability, ICC(1, 1), is also included from the CCBD dataset. ^†^FreeSurfer labels in version 6.0; labels are named slightly different in version 5.3. ICC values between .75 and 1.00 are considered ‘excellent,’ .60–.74 is ‘good,’ .40–.59 is ‘fair,’ and below .40 is ‘poor’ (Cicchetti, 1994).

| Sulci Name | FreeSurfer Label† | Hemi. | OASIS | DLBS | SALD | CCBD |  |
| --- | --- | --- | --- | --- | --- | --- | --- |
|  |  |  | *r(Age)* | *r(Age)* | *r(Age)* | *ICC(1,1)* | 95% CI of ICC |
| Central | S_central | L | -.517 | -.205 | -.346 | .848 | [ 0.772, 0.912] |
|  |  | R | -.505 | -.256 | -.348 | .860 | [ 0.789, 0.919] |
| Post-central | S_postcentral | L | -.371 | -.264 | -.268 | .965 | [ 0.944, 0.981] |
|  |  | R | -.436 | -.246 | -.330 | .890 | [ 0.831, 0.937] |
| Superior Frontal | S_front_sup | L | -.523 | -.454 | -.397 | .899 | [ 0.844, 0.943] |
|  |  | R | -.413 | -.465 | -.444 | .886 | [ 0.825, 0.935] |
| Inferior Frontal | S_front_inf | L | -.517 | -.490 | -.491 | .932 | [ 0.893, 0.962] |
|  |  | R | -.496 | -.480 | -.490 | .915 | [ 0.868, 0.952] |
| Parieto-occipital | S_parieto_occipital | L | -.145 | -.093 | -.241 | .979 | [ 0.966, 0.989] |
|  |  | R | -.124 | .059 | -.229 | .970 | [ 0.952, 0.984] |
| Occipito-temporal | S_oc-temp_med&Lingual | L | -.509 | -.323 | -.263 | .953 | [ 0.926, 0.974] |
|  |  | R | -.404 | -.316 | -.281 | .913 | [ 0.864, 0.951] |
| Middle occipital and lunate | S_oc_middle&Lunatus | L | -.290 | -.167 | -.150 | .949 | [ 0.919, 0.972] |
|  |  | R | -.288 | -.120 | -.132 | .922 | [ 0.879, 0.956] |
| Marginal part of cingulate | S_cingul-Marginalis | L | -.092 | -.035 | -.268 | .952 | [ 0.925, 0.974] |
|  |  | R | -.032 | -.017 | -.156 | .918 | [ 0.872, 0.954] |
| Mean |  |  | -.465 | -.645 | -.600 | .972 | [ 0.955, 0.985] |

Age-related relationships for each sulcus were relatively consistent between the two Western lifespan datasets (OASIS and DLBS), but age-related differences in sulcal width (but not depth) were markedly weaker in the East Asian lifespan dataset (SALD). This finding will need to be studied further, but may be related to gross differences in anatomical structure (Kochunov et al., 2003; Longstreth et al., 2000; Tang et al., 2010)–and motivates the need to aging in samples that vary in ethnicity/race and are otherwise not of a so-called WEIRD (Western, Educated, Industrialized, Rich, and Democratic) demographic (Madan, 2017). Additionally, there did not appear to be a significant influence of field strength (i.e., 1.5 T for the OASIS dataset vs. 3 T for the DLBS dataset) on estimates of sulcal morphology. Importantly, test-retest reliability, *ICC*(1, 1), was particularly good for the sulcal depth across individual sulci.

To obtain a coarse summary measure across sulci, we averaged the sulcal width across the 16 individual sulci for each individual, and with each dataset, and examined the relationship between mean sulcal width with age. These correlations, shown in Table 1, indicate that the mean sulcal width was generally a better indicator of age-related differences in sulcal morphology than individual sulci, and had increased test-retest reliability. Mean sulcal depth was similarly more sensitive to age-related differences than for an individual sulcus (e.g., it is unclear why the relationship between age and width of the central sulcus differed between samples) and the magnitude of this relationship was more consistent across datasets. Reliability was even higher for mean sulcal depth than mean sulcal width.

### 4.2 Comparison with other age-related structural differences

Within each dataset, mean sulcal depth and width correlated with age, as shown in Tables 1 and 2. Of course, other measures of brain morphology also differ with age, such as mean (global) cortical thickness [OASIS: *r*(308) = *−.*793, *p* < .001; DLBS: *r*(310) = *−.*759, *p* < .001; SALD: *r*(479) = *−.*642, *p* < .001]. and volume of the third ventricle (ICV-corrected) [OASIS: *r*(308) = .665, *p* < .001; DLBS: *r*(310) = .677, *p* < .001; SALD: *r*(479) = .328, *p* < .001]. Previous studies have demonstrated that both of these measures are robust estimates of age-related differences in brain structure (Fjell et al., 2009; Hogstrom et al., 2013; Hutton et al., 2009; Lemaitre et al., 2012; Madan & Kensinger, 2016, 2017a; Madan, 2018; McKay et al., 2014; Salat et al., 2004; Sowell et al., 2003, 2007; Walhovd et al., 2011).

To test if these mean sulcal measures served as distinct measures of age-related differences in brain morphology, beyond those provided by other measures, such as mean cortical thickness and volume of the third ventricle, we conducted partial correlations that controlled for these two other measures of age-related atrophy. Mean sulcal width [OASIS: *r_p_*(306) = .188, *p* < .001; DLBS: *r_p_*(308) = .177, *p* = .002; SALD: *r*(477) = .003, *p* = .96] and depth [OASIS: *r_p_*(306) = *−.*443, *p* < .001; DLBS: *r_p_*(308) = *−.*397, *p* < .001; SALD: *r_p_*(477) = *−.*534, *p* < .001] both explained unique variance in relation to age. Thus, even though more established measures of age-related differences in brain morphology were replicated here, the additional sulcal measures captured aspects of aging that are not accounted for by these extant measures, indicating that these sulcal measures are worth pursuing further and are not redundant with other measures of brain structure. Providing additional support for this, mean sulcal width and depth were only weakly related to each other [OASIS: *r*(308) = *−.*192, *p* < .001; DLBS: *r*(310) = .092, *p* = .104; SALD: *r*(479) = .119, *p* = .009].

As with the individual sulci measures, we did observe a difference between samples where some age-related measures were less sensitive in the East Asian lifespan sample (SALD), here in the ventricle volume correlation and the unsurprisingly weaker age relationship in the partial correlation using sulcal width. These sample differences are puzzling, though there is a general correspondence between the two Western samples. Given that much of the literature is also based on Western samples, we think further research with East Asian samples, and particularly comparing samples with the same analysis pipeline, is necessary to shed further light on this initial finding.

## 5 Conclusion

Differences in sulcal width and depth are quite visually prominent, but are not often quantified when examining individual differences in cortical structure. Here we examined age-related differences in both sulcal measures as a proof-of-principle to demonstrate the utility of the calcSulc toolbox that accompanies this paper and is designed to closely compliments the standard FreeSurfer pipeline. This allows for the additional measurement of sulcal morphology, to add to the extant measures of brain morphology such as cortical thickness, area, and gyrification. Critically, this approach uses the same landmarks and boundaries as in the Destrieux et al. (2010) parcellation atlas, in contrast to all previous approaches to characterize sulcal features. This toolbox is now made freely available as supplemental to this paper: https://cmadan.github.io/calcSulc/.

Using this approach, here we demonstrate age-related differences in sulcal width and depth, as well as high test-retest reliability. Since individual differences in sulcal morphology are sufficiently distinct from those characterized by other brain morphology measures, this approach should complement extant work of investigating factors that influence brain morphology, e.g., see Figure 3 of Madan and Kensinger (2018). Given the flexibility in the methodological approach, these measures can be readily applied to other samples after being initially processed with FreeSurfer.

## Acknowledgments

MRI data used in the preparation of this article were obtained from several sources, data were provided in part by: (1) the Open Access Series of Imaging Studies (OASIS) (Marcus et al., 2007); (2) wave 1 of the Dallas Lifespan Brain Study (DLBS) led by Dr. Denise Park and distributed through INDI (Mennes et al., 2013) and NITRC (Kennedy et al., 2016); (3) the Southwest University Adult Lifespan Dataset (SALD) (Wei et al., 2018), also made available through INDI and hosted on NITRC; and (4) the Center for Cognition and Brain Disorders (CCBD) (Chen et al., 2015) as dataset HNU1 in the Consortium for Reliability and Reproducibility (CoRR) (Zuo et al., 2014).

## Footnotes

1 Note that analyses reported in these previous papers were based on preprocessing in FreeSurfer 5.3.0, rather than FreeSurfer 6.0.

## References

Andersen, S. K., Jakobsen, C. E., Pedersen, C. H., Rasmussen, A. M., Plocharski, M., & Østergaard, L. R. (2015). Classification of Alzheimer’s disease from MRI using sulcal morphology. In Scandinavian Conference on Image Analysis (SCIA): Image Analysis (pp. 103–113). Springer. doi: 10.1007/978-3-319-19665-7_9

Andreasen, N. C., Harris, G., Cizadlo, T., Arndt, S., O’Leary, D. S., Swayze, V., & Flaum, M. (1994). Techniques for measuring sulcal/gyral patterns in the brain as visualized through magnetic resonance scanning: BRAINPLOT and BRAINMAP. Proceedings of the National Academy of Sciences, 91, 93–97. doi: 10.1073/pnas.91.1.93

Asendorpf, J., & Wallbott, H. G. (1979). Maße der Beobachterübereinstimmung: ein systematischer Vergleich. Zeitschrift für Sozialpsychologie, 10, 243–252.

Auzias, G., Brun, L., Deruelle, C., & Coulon, O. (2015). Deep sulcal landmarks: Algorithmic and conceptual improvements in the definition and extraction of sulcal pits. NeuroImage, 111, 12–25. doi: 10.1016/j.neuroimage.2015.02.008

Bartko, J. J. (1966). The intraclass correlation coefficient as a measure of reliability. Psychological Reports, 19, 3–11. doi: 10.2466/pr0.1966.19.1.3

Beeston, C. J., & Taylor, C. J. (2000). Automatic landmarking of cortical sulci. In Medical image computing and computer-assisted intervention–MICCAI 2000 (pp. 125–133). Springer Berlin Heidelberg. doi: 10.1007/978-3-540-40899-4_13

Behnke, K. J., Rettmann, M. E., Pham, D. L., Shen, D., Resnick, S. M., Davatzikos, C., & Prince, J. L. (2003). Automatic classification of sulcal regions of the human brain cortex using pattern recognition. In M. Sonka & J. M. Fitzpatrick (Eds.), Medical imaging 2003: Image processing (pp. 1499–1510). SPIE. doi: 10.1117/12.480834

Cai, K., Xu, H., Guan, H., Zhu, W., Jiang, J., Cui, Y., … Wen, W. (2017). Identification of early-stage Alzheimer’s disease using sulcal morphology and other common neuroimaging indices. PLOS ONE, 12, e0170875. doi: 10.1371/journal.pone.0170875

Campero, A., Ajler, P., Emmerich, J., Goldschmidt, E., Martins, C., & Rhoton, A. (2014). Brain sulci and gyri: A practical anatomical review. Journal of Clinical Neuroscience, 21, 2219–2225. doi: 10.1016/j.jocn.2014.02.024

Cao, B., Mwangi, B., Passos, I. C., Wu, M.-J., Keser, Z., Zunta-Soares, G. B., … Soares, J. C. (2017). Lifespan gyrification trajectories of human brain in healthy individuals and patients with major psychiatric disorders. Scientific Reports, 7, 511. doi: 10.1038/s41598-017-00582-1

Chan, M. Y., Park, D. C., Savalia, N. K., Petersen, S. E., & Wig, G. S. (2014). Decreased segregation of brain systems across the healthy adult lifespan. Proceedings of the National Academy of Sciences USA, 111, E4997–E5006. doi: 10.1073/pnas.1415122111

Chen, B., Xu, T., Zhou, C., Wang, L., Yang, N., Wang, Z., … Weng, X.-C. (2015). Individual variability and test-retest reliability revealed by ten repeated resting-state brain scans over one month. PLOS ONE, 10, e0144963. doi: 10.1371/journal.pone.0144963

Chen, G., Taylor, P. A., Haller, S. P., Kircanski, K., Stoddard, J., Pine, D. S., … Cox, R. W. (2018). Intraclass correlation: Improved modeling approaches and applications for neuroimaging. Human Brain Mapping, 39, 1187–1206. doi: 10.1002/hbm.23909

Cicchetti, D. V. (1994). Guidelines, criteria, and rules of thumb for evaluating normed and standardized assessment instruments in psychology. Psychological Assessment, 6, 284–290. doi: 10.1037/1040-3590.6.4.284

Coffey, C. E., Wilkinson, W. E., Parashos, L., Soady, S., Sullivan, R. J., Patterson, L. J., … Djang, W. T. (1992). Quantitative cerebral anatomy of the aging human brain: A cross-sectional study using magnetic resonance imaging. Neurology, 42, 527–527. doi: 10.1212/wnl.42.3.527

Destrieux, C., Fischl, B., Dale, A., & Halgren, E. (2010). Automatic parcellation of human cortical gyri and sulci using standard anatomical nomenclature. NeuroImage, 53, 1–15. doi: 10.1016/j.neuroimage.2010.06.010

Drayer, B. P. (1988). Imaging of the aging brain. Part I. normal findings. Radiology, 166, 785–796. doi: 10.1148/radiology.166.3.3277247

Elkis, H., Friedman, L., Wise, A., & Meltzer, H. Y. (1995). Meta-analyses of studies of ventricular enlargement and cortical sulcal prominence in mood disorders. Archives of General Psychiatry, 52, 735–746. doi: 10.1001/archpsyc.1995.03950210029008

Eskildsen, S. F., Uldahl, M., & Ostergaard, L. R. (2005). Extraction of the cerebral cortical boundaries from MRI for measurement of cortical thickness. In J. M. Fitzpatrick & J. M. Reinhardt (Eds.), Medical imaging 2005: Image processing. SPIE. doi: 10.1117/12.595145

Fischl, B. (2012). FreeSurfer. NeuroImage, 62, 774–781. doi: 10.1016/j.neuroimage.2012.01.021

Fischl, B., & Dale, A. M. (2000). Measuring the thickness of the human cerebral cortex from magnetic resonance images. Proceedings of the National Academy of Sciences USA, 97, 11050–11055. doi: 10.1073/pnas.200033797

Fjell, A. M., Westlye, L. T., Amlien, I., Espeseth, T., Reinvang, I., Raz, N., … Walhovd, K. B. (2009). High consistency of regional cortical thinning in aging across multiple samples. Cerebral Cortex, 19, 2001–2012. doi: 10.1093/cercor/bhn232

Hallgren, K. A. (2012). Computing inter-rater reliability for observational data: An overview and tutorial. Tutorials in Quantitative Methods for Psychology, 8, 23–34. doi: 10.20982/tqmp.08.1.p023

Hamelin, L., Bertoux, M., Bottlaender, M., Corne, H., Lagarde, J., Hahn, V., … Sarazin, M. (2015). Sulcal morphology as a new imaging marker for the diagnosis of early onset Alzheimer’s disease. Neurobiology of Aging, 36, 2932–2939. doi: 10.1016/j.neurobiolaging.2015.04.019

Hogstrom, L. J., Westlye, L. T., Walhovd, K. B., & Fjell, A. M. (2013). The structure of the cerebral cortex across adult life: Age-related patterns of surface area, thickness, and gyrification. Cerebral Cortex, 23, 2521–2530. doi: 10.1093/cercor/bhs231

Huckman, M. S., Fox, J., & Topel, J. (1975). The validity of criteria for the evaluation of cerebral atrophy by computed tomography. Radiology, 116, 85–92. doi: 10.1148/116.1.85

Hutton, C., Draganski, B., Ashburner, J., & Weiskopf, N. (2009). A comparison between voxel-based cortical thickness and voxel-based morphometry in normal aging. NeuroImage, 48, 371–380. doi: 10.1016/j.neuroimage.2009.06.043

Im, K., Jo, H. J., Mangin, J.-F., Evans, A. C., Kim, S. I., & Lee, J.-M. (2010). Spatial distribution of deep sulcal landmarks and hemispherical asymmetry on the cortical surface. Cerebral Cortex, 20, 602–611. doi: 10.1093/cercor/bhp127

Jacoby, R. J., Levy, R., & Dawson, J. M. (1980). Computed tomography in the elderly: I. The normal population. British Journal of Psychiatry, 136, 249–255. doi: 10.1192/bjp.136.3.249

John, J. P., Wang, L., Moffitt, A. J., Singh, H. K., Gado, M. H., & Csernansky, J. G. (2006). Inter-rater reliability of manual segmentation of the superior, inferior and middle frontal gyri. Psychiatry Research: Neuroimaging, 148, 151–163. doi: 10.1016/j.pscychresns.2006.05.006

Jones, S. E., Buchbinder, B. R., & Aharon, I. (2000). Three-dimensional mapping of cortical thickness using Laplace’s equation. Human Brain Mapping, 11, 12–32. doi: 10.1002/1097-0193(200009)11:1<12::aid-hbm20>3.0.co;2-k

Kennedy, D. N., Haselgrove, C., Riehl, J., Preuss, N., & Buccigrossi, R. (2016). The NITRC image repository. NeuroImage, 124, 1069–1073. doi: 10.1016/j.neuroimage.2015.05.074

Kennedy, K. M., Rodrigue, K. M., Bischof, G. N., Hebrank, A. C., Reuter-Lorenz, P. A., & Park, D. C. (2015). Age trajectories of functional activation under conditions of low and high processing demands: An adult lifespan fMRI study of the aging brain. NeuroImage, 104, 21–34. doi: 10.1016/j.neuroimage.2014.09.056

Kippenhan, J. S., Olsen, R. K., Mervis, C. B., Morris, C. A., Kohn, P., Meyer-Lindenberg, A., & Berman, K. F. (2005). Genetic contributions to human gyrification: Sulcal morphometry in Williams syndrome. Journal of Neuroscience, 25, 7840–7846. doi: 10.1523/jneurosci.1722-05.2005

Kochunov, P., Fox, P., Lancaster, J., Tan, L. H., Amunts, K., Zilles, K., … Gao, J. H. (2003). Localized morphological brain differences between english-speaking caucasians and chinese-speaking asians: new evidence of anatomical plasticity. NeuroReport, 14, 961–964. doi: 10.1097/01.wnr.0000075417.59944.00

Kochunov, P., Mangin, J.-F., Coyle, T., Lancaster, J., Thompson, P., Rivière, D., … Fox, P. T. (2005). Age-related morphology trends of cortical sulci. Human Brain Mapping, 26, 210–220. doi: 10.1002/hbm.20198

Kochunov, P., Rogers, W., Mangin, J.-F., & Lancaster, J. (2012). A library of cortical morphology analysis tools to study development, aging and genetics of cerebral cortex. Neuroinformatics, 10, 81–96. doi: 10.1007/s12021-011-9127-9

Kochunov, P., Thompson, P. M., Coyle, T. R., Lancaster, J. L., Kochunov, V., Royall, D., … Fox, P. T. (2008). Relationship among neuroimaging indices of cerebral health during normal aging. Human Brain Mapping, 29, 36–45. doi: 10.1002/hbm.20369

Koo, T. K., & Li, M. Y. (2016). A guideline of selecting and reporting intraclass correlation coefficients for reliability research. Journal of Chiropractic Medicine, 15, 155–163. doi: 10.1016/j.jcm.2016.02.012

Laffey, P. A., Peyster, R. G., Nathan, R., Haskin, M. E., & McGinley, J. A. (1984). Computed tomography and aging: Results in a normal elderly population. Neuroradiology, 26, 273–278. doi: 10.1007/BF00339770

Lamont, A. J., Mortby, M. E., Anstey, K. J., Sachdev, P. S., & Cherbuin, N. (2014). Using sulcal and gyral measures of brain structure to investigate benefits of an active lifestyle. NeuroImage, 91, 353–359. doi: 10.1016/j.neuroimage.2014.01.008

Largen, J. W., Smith, R. C., Calderon, M., Baumgartner, R., Lu, R. B., Schoolar, J. C., & Ravichandran, G. K. (1984). Abnormalities of brain structure and density in schizophrenia. Biological Psychiatry, 19, 991–1013.

Le Goualher, G., Barillot, C., Bizais, Y. J., & Scarabin, J.-M. (1996). Three-dimensional segmentation of cortical sulci using active models. In M. H. Loew & K. M. Hanson (Eds.), Medical imaging 1996: Image processing (pp. 254–263). SPIE. doi: 10.1117/12.237928

Le Goualher, G., Collins, D. L., Barillot, C., & Evans, A. C. (1998). Automatic identificaiton of cortical sulci using a 3d probabilistic atlas. In Medical image computing and computer-assisted intervention – MICCAI’98 (pp. 509–518). Springer Berlin Heidelberg. doi: 10.1007/bfb0056236

Le Troter, A., Auzias, G., & Coulon, O. (2012). Automatic sulcal line extraction on cortical surfaces using geodesic path density maps. NeuroImage, 61, 941–949. doi: 10.1016/j.neuroimage.2012.04.021

Lee, J. K., Lee, J.-M., Kim, J. S., Kim, I. Y., Evans, A. C., & Kim, S. I. (2006). A novel quantitative cross-validation of different cortical surface reconstruction algorithms using MRI phantom. NeuroImage, 31, 572–584. doi: 10.1016/j.neuroimage.2005.12.044

Lemaitre, H., Goldman, A. L., Sambataro, F., Verchinski, B. A., Meyer-Lindenberg, A., Weinberger, D. R., & Mattay, V. S. (2012). Normal age-related brain morphometric changes: nonuniformity across cortical thickness, surface area and gray matter volume? Neurobiology of Aging, 33, 617.e1–617.e9. doi: 10.1016/j.neurobiolaging.2010.07.013

Leong, R. L., Lo, J. C., Sim, S. K., Zheng, H., Tandi, J., Zhou, J., & Chee, M. W. (2017). Longitudinal brain structure and cognitive changes over 8 years in an east asian cohort. NeuroImage, 147, 852–860. doi: 10.1016/j.neuroimage.2016.10.016

Li, G., Guo, L., Nie, J., & Liu, T. (2010). An automated pipeline for cortical sulcal fundi extraction. Medical Image Analysis, 14, 343–359. doi: 10.1016/j.media.2010.01.005

Li, G., & Shen, D. (2011). Consistent sulcal parcellation of longitudinal cortical surfaces. NeuroImage, 57, 76–88. doi: 10.1016/j.neuroimage.2011.03.064

Liu, T., Lipnicki, D. M., Zhu, W., Tao, D., Zhang, C., Cui, Y., … Wen, W. (2012). Cortical gyrification and sulcal spans in early stage Alzheimer’s disease. PLOS ONE, 7, e31083. doi: 10.1371/journal.pone.0031083

Liu, T., Sachdev, P. S., Lipnicki, D. M., Jiang, J., Geng, G., Zhu, W., … Wen, W. (2013). Limited relationships between two-year changes in sulcal morphology and other common neuroimaging indices in the elderly. NeuroImage, 83, 12–17. doi: 10.1016/j.neuroimage.2013.06.058

Liu, T., Wen, W., Zhu, W., Kochan, N. A., Trollor, J. N., Reppermund, S., … Sachdev, P. S. (2011). The relationship between cortical sulcal variability and cognitive performance in the elderly. NeuroImage, 56, 865–873. doi: 10.1016/j.neuroimage.2011.03.015

Liu, T., Wen, W., Zhu, W., Trollor, J., Reppermund, S., Crawford, J., … Sachdev, P. (2010). The effects of age and sex on cortical sulci in the elderly. NeuroImage, 51, 19–27. doi: 10.1016/j.neuroimage.2010.02.016

Lohmann, G., & von Cramon, D. Y. (2000). Automatic labelling of the human cortical surface using sulcal basins. Medical Image Analysis, 4, 179–188. doi: 10.1016/s1361-8415(00)00024-4

Lohmann, G., von Cramon, D. Y., & Colchester, A. C. F. (2008). Deep sulcal landmarks provide an organizing framework for human cortical folding. Cerebral Cortex, 18, 1415–1420. doi: 10.1093/cercor/bhm174

Longstreth, W. T., Arnold, A. M., Manolio, T. A., Burke, G. L., Bryan, N., Jungreis, C. A., … Fried, L. (2000). Clinical correlates of ventricular and sulcal size on cranial magnetic resonance imaging of 3,301 elderly people. Neuroepidemiology, 19, 30–42. doi: 10.1159/000026235

Madan, C. R. (2017). Advances in studying brain morphology: The benefits of open-access data. Frontiers in Human Neuroscience, 11, 405. doi: 10.3389/fnhum.2017.00405

Madan, C. R. (2018). Age differences in head motion and estimates of cortical morphology. PeerJ, 6, e5176. doi: 10.7717/peerj.5176

Madan, C. R. (2019). Shape-related characteristics of age-related differences in subcortical structures. Aging & Mental Health, 23, 800–810. doi: 10.1080/13607863.2017.1421613

Madan, C. R., & Kensinger, E. A. (2016). Cortical complexity as a measure of age-related brain atrophy. NeuroImage, 134, 617–629. doi: 10.1016/j.neuroimage.2016.04.029

Madan, C. R., & Kensinger, E. A. (2017a). Age-related differences in the structural complexity of subcortical and ventricular structures. Neurobiology of Aging, 50, 87–95. doi: 10.1016/j.neurobiolaging.2016.10.023

Madan, C. R., & Kensinger, E. A. (2017b). Test–retest reliability of brain morphology estimates. Brain Informatics, 4, 107–121. doi: 10.1007/s40708-016-0060-4

Madan, C. R., & Kensinger, E. A. (2018). Predicting age from cortical structure across the lifespan. European Journal of Neuroscience, 47, 399–416. doi: 10.1111/ejn.13835

Mangin, J.-F., Riviere, D., Cachia, A., Duchesnay, E., Cointepas, Y., Papadopoulos-Orfanos, D., … Regis, J. (2004). Object-based morphometry of the cerebral cortex. IEEE Transactions on Medical Imaging, 23, 968–982. doi: 10.1109/tmi.2004.831204

Mangin, J.-F., Rivière, D., Coulon, O., Poupon, C., Cachia, A., Cointepas, Y., … Papadopoulos-Orfanos, D. (2004). Coordinate-based versus structural approaches to brain image analysis. Artificial Intelligence in Medicine, 30, 177–197. doi: 10.1016/s0933-3657(03)00064-2

Marcus, D. S., Wang, T. H., Parker, J., Csernansky, J. G., Morris, J. C., & Buckner, R. L. (2007). Open Access Series of Imaging Studies (OASIS): Cross-sectional MRI data in young, middle aged, nondemented, and demented older adults. Journal of Cognitive Neuroscience, 19, 1498–1507. doi: 10.1162/jocn.2007.19.9.1498

McGraw, K. O., & Wong, S. P. (1996). Forming inferences about some intraclass correlation coefficients. Psychological Methods, 1, 30–46. doi: 10.1037/1082-989x.1.1.30

McKay, D. R., Knowles, E. E. M., Winkler, A. A. M., Sprooten, E., Kochunov, P., Olvera, R. L., … Glahn, D. C. (2014). Influence of age, sex and genetic factors on the human brain. Brain Imaging and Behavior, 8, 143–152. doi: 10.1007/s11682-013-9277-5

Mennes, M., Biswal, B. B., Castellanos, F. X., & Milham, M. P. (2013). Making data sharing work: The FCP/INDI experience. NeuroImage, 82, 683–691. doi: 10.1016/j.neuroimage.2012.10.064

Mikhael, S., Hoogendoorn, C., Valdes-Hernandez, M., & Pernet, C. (2018). A critical analysis of neuroanatomical software protocols reveals clinically relevant differences in parcellation schemes. NeuroImage, 170, 348–364. doi: 10.1016/j.neuroimage.2017.02.082

Ming, J., Harms, M. P., Morris, J. C., Beg, M. F., & Wang, L. (2015). Integrated cortical structural marker for Alzheimer’s disease. Neurobiology of Aging, 36, S53–S59. doi: 10.1016/j.neurobiolaging.2014.03.042

Nowinski, W. L., Raphel, J. K., & Nguyen, B. T. (1996). Atlas-based identification of cortical sulci. In Y. Kim (Ed.), Medical imaging 1996: Image display (pp. 64–74). SPIE. doi: 10.1117/12.238488

Oguz, I., Cates, J., Fletcher, T., Whitaker, R., Cool, D., Aylward, S., & Styner, M. (2008). Cortical correspondence using entropy-based particle systems and local features. In 2008 5th IEEE International Symposium on Biomedical Imaging: From nano to macro (pp. 1637–1640). IEEE. doi: 10.1109/isbi.2008.4541327

Ono, M., Kubick, S., & Abernathey, C. D. (1990). Atlas of the cerebral sulci. Thieme.

Palaniyappan, L., Park, B., Balain, V., Dangi, R., & Liddle, P. (2015). Abnormalities in structural covariance of cortical gyrification in schizophrenia. Brain Structure and Function, 220, 2059–2071. doi: 10.1007/s00429-014-0772-2

Pasquier, F., Leys, D., Weerts, J. G., Mounier-Vehier, F., Barkhof, F., & Scheltens, P. (1996). Inter- and intraobserver reproducibility of cerebral atrophy assessment on MRI scans with hemispheric infarcts. European Neurology, 36, 268–272. doi: 10.1159/000117270

Perrot, M., Rivière, D., & Mangin, J.-F. (2011). Cortical sulci recognition and spatial normalization. Medical Image Analysis, 15, 529–550. doi: 10.1016/j.media.2011.02.008

Pizzagalli, F., Auzias, G., Kochunov, P., Faskowitz, J. I., Thompson, P. M., & Jahanshad, N. (2017). The core genetic network underlying sulcal morphometry. In E. Romero, N. Lepore, J. Brieva, & I. Larrabide (Eds.), International Symposium on Medical Information Processing and Analysis. SPIE. doi: 10.1117/12.2256959

Plocharski, M., & Østergaard, L. R. (2016). Extraction of sulcal medial surface and classification of Alzheimer’s disease using sulcal features. Computer Methods and Programs in Biomedicine, 133, 35–44. doi: 10.1016/j.cmpb.2016.05.009

Rajaratnam, N. (1960). Reliability formulas for independent decision data when reliability data are matched. Psychometrika, 25, 261–271. doi: 10.1007/bf02289730

Reiner, P., Jouvent, E., Duchesnay, E., Cuingnet, R., Mangin, J.-F., & Chabriat, H. (2012). Sulcal span in Alzheimer’s disease, amnestic mild cognitive impairment, and healthy controls. Journal of Alzheimer’s Disease, 29, 605–613. doi: 10.3233/JAD-2012-111622

Rhoton, A. L. (2007). The cerebrum. Neurosurgery, 61, SHC-37–SHC-119. doi: 10.1227/01.neu.0000255490.88321.ce

Rieder, R. O., Donnelly, E. F., Herdt, J. R., & Waldman, I. N. (1979). Sulcal prominence in young chronic schizophrenic patients: CT scan findings associated with impairment on neuropsychological tests. Psychiatry Research, 1, 1–8. doi: 10.1016/0165-1781(79)90021-0

Rivière, D., Mangin, J.-F., Papadopoulos-Orfanos, D., Martinez, J.-M., Frouin, V., & Régis, J. (2002). Automatic recognition of cortical sulci of the human brain using a congregation of neural networks. Medical Image Analysis, 6, 77–92. doi: 10.1016/s1361-8415(02)00052-x

Royackkers, N., Desvignes, M., Fawal, H., & Revenu, M. (1999). Detection and statistical analysis of human cortical sulci. NeuroImage, 10, 625–641. doi: 10.1006/nimg.1999.0512

Salat, D. H., Buckner, R. L., Snyder, A. Z., Greve, D. N., Desikan, R. S. R., Busa, E., … Fischl, B. (2004). Thinning of the cerebral cortex in aging. Cerebral Cortex, 14, 721–730. doi: 10.1093/cercor/bhh032

Schaer, M., Cuadra, M. B., Schmansky, N., Fischl, B., Thiran, J.-P., & Eliez, S. (2012). How to measure cortical folding from MR images: A step-by-step tutorial to compute local gyrification index. Journal of Visualized Experiments, e3417. doi: 10.3791/3417

Scheltens, P., Pasquier, F., Weerts, J. G., Barkhof, F., & Leys, D. (1997). Qualitative assessment of cerebral atrophy on MRI: Inter- and intra-observer reproducibility in dementia and normal aging. European Neurology, 37, 95–99. doi: 10.1159/000117417

Shrout, P. E., & Fleiss, J. L. (1979). Intraclass correlations: Uses in assessing rater reliability. Psychological Bulletin, 86, 420–428. doi: 10.1037/0033-2909.86.2.420

Sowell, E. R., Peterson, B. S., Kan, E., Woods, R. P., Yoshii, J., Bansal, R., … Toga, A. W. (2007). Sex differences in cortical thickness mapped in 176 healthy individuals between 7 and 87 years of age. Cerebral Cortex, 17, 1550–1560. doi: 10.1093/cercor/bhl066

Sowell, E. R., Peterson, B. S., Thompson, P. M., Welcome, S. E., Henkenius, A. L., & Toga, A. W. (2003). Mapping cortical change across the human life span. Nature Neuroscience, 6, 309–315. doi: 10.1038/nn1008

Tang, Y., Hojatkashani, C., Dinov, I. D., Sun, B., Fan, L., Lin, X., … Toga, A. W. (2010). The construction of a chinese MRI brain atlas: A morphometric comparison study between chinese and caucasian cohorts. NeuroImage, 51, 33–41. doi: 10.1016/j.neuroimage.2010.01.111

ten Donkelaar, H. J., Tzourio-Mazoyer, N., & Mai, J. K. (2018). Toward a common terminology for the gyri and sulci of the human cerebral cortex. Frontiers in Neuroanatomy, 12, 93. doi: 10.3389/fnana.2018.00093

Thompson, P. M., Schwartz, C., Lin, R. T., Khan, A. A., & Toga, A. W. (1996). Three-dimensional statistical analysis of sulcal variability in the human brain. Journal of Neuroscience, 16, 4261–4274. doi: 10.1523/jneurosci.16-13-04261.1996

Tomlinson, B., Blessed, G., & Roth, M. (1968). Observations on the brains of non-demented old people. Journal of the Neurological Sciences, 7, 331–356. doi: 10.1016/0022-510x(68)90154-8

Vaillant, M., & Davatzikos, C. (1997). Finding parametric representations of the cortical sulci using an active contour model. Medical Image Analysis, 1, 295–315. doi: 10.1016/s1361-8415(97)85003-7

Walhovd, K. B., Westlye, L. T., Amlien, I., Espeseth, T., Reinvang, I., Raz, N., … Fjell, A. M. (2011). Consistent neuroanatomical age-related volume differences across multiple samples. Neurobiology of Aging, 32, 916–932. doi: 10.1016/j.neurobiolaging.2009.05.013

Wei, D., Zhuang, K., Ai, L., Chen, Q., Yang, W., Liu, W., … Qiu, J. (2018). Structural and functional brain scans from the cross-sectional southwest university adult lifespan dataset. Scientific Data, 5, 180134. doi: 10.1038/sdata.2018.134

Welker, W. (1990). Why does cerebral cortex fissure and fold? In E. G. Jones & A. Peters (Eds.), Cerebral cortex (Vol. 8B, pp. 3–136). Springer US. doi: 10.1007/978-1-4615-3824-0_1

Yang, F., & Kruggel, F. (2008). Automatic segmentation of human brain sulci. Medical Image Analysis, 12, 442–451. doi: 10.1016/j.media.2008.01.003

Yue, N. C., Arnold, A. M., Longstreth, W. T., Elster, A. D., Jungreis, C. A., O’Leary, D. H., … Bryan, R. N. (1997). Sulcal, ventricular, and white matter changes at MR imaging in the aging brain: Data from the cardiovascular health study. Radiology, 202, 33–39. doi: 10.1148/radiology.202.1.8988189

Yun, H. J., Im, K., Yang, J.-J., Yoon, U., & Lee, J.-M. (2013). Automated sulcal depth measurement on cortical surface reflecting geometrical properties of sulci. PLOS ONE, 8, e55977. doi: 10.1371/journal.pone.0055977

Zuo, X.-N., Anderson, J. S., Bellec, P., Birn, R. M., Biswal, B. B., Blautzik, J., … Milham, M. P. (2014). An open science resource for establishing reliability and reproducibility in functional connectomics. Scientific Data, 1, 140049. doi: 10.1038/sdata.2014.49

